# Drosophila’s sensory responses to bacterial peptidoglycan integrates positive and negative signals

**DOI:** 10.1101/2024.12.01.626038

**Authors:** Martina Montanari, Gérard Manière, Ambra Masuzzo, Romane Milleville, Yaël Grosjean, C. Léopold Kurz, Julien Royet

## Abstract

Interactions between animals, including humans, and surrounding microbes are governed by a delicate balance, crucial for survival. Animals must distinguish and respond adequately to beneficial and harmful microbes to maintain homeostasis. Recent research suggests that bacterial components such as lipopolysaccharide and peptidoglycan (PGN) influence host behavior by modulating neuronal activity. PGN detection by specific neurons can prompt infected female flies to reduce oviposition or trigger avoidance behaviors via gustatory neurons. Using behavioral assays and calcium imaging, we found that PGNs can also act as attractants, activating the sweet taste circuit in a concentration-dependent manner. Our findings demonstrate that flies integrate PGN-derived positive and negative signals to make ad hoc decisions. This dual response underlines the need for *Drosophila* to distinguish between different concentrations of compounds in their environment, integrating sensory data to navigate efficiently in microbe-co-inhabited environments.

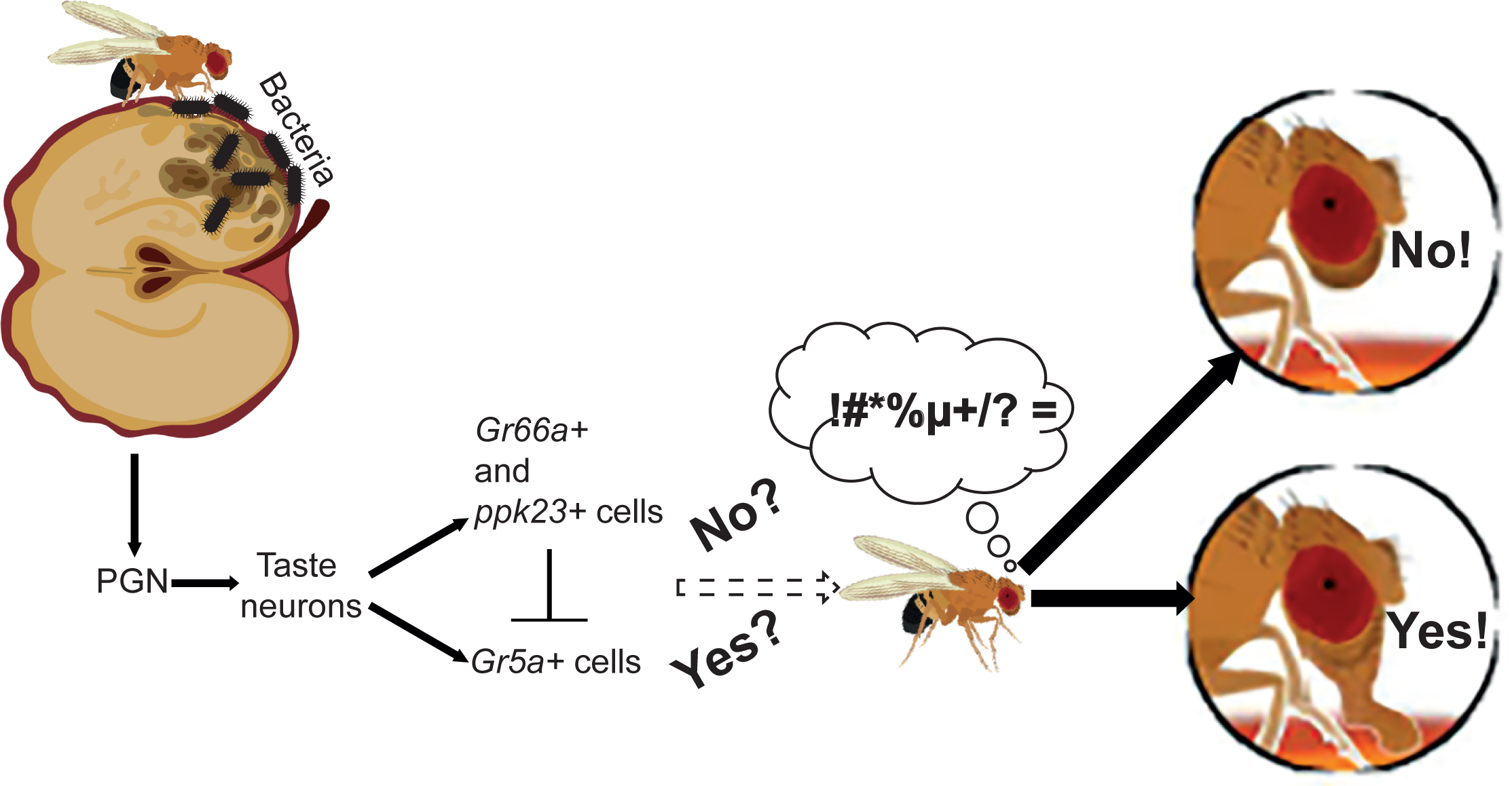

**Highlights:**

- Bacterial PGN is attractive to flies
- Gr5a sweet gustatory neurons are activated by PGN
- Adult PER to PGN is not directly influenced by larval life
- Fly gustatory response to PGN integrates both attractive and aversive signals

## Introduction

The interactions between animals, including humans, and the microbes surrounding them are governed by a subtle balance. To understand the mechanisms underlying this equilibrium, the insect *Drosophila melanogaster* serves as a powerful model, as microorganisms are essential for the host’s development, survival, nutrients, or making certain food sources digestible (Storelli et al., 2011). Some beneficial microbes occupy ecological niches that would otherwise be filled by harmful microorganisms if they were absent (Kim et al., 2010). Conversely, certain pathogenic species must be detected, avoided, or even actively combated by animals (Charroux et al., 2020; Chen et al., 2024; Tleiss et al., 2024). The ability of animals to detect, identify, and respond appropriately to the microbial world is therefore essential for their homeostasis and sustainability.

Recent studies have begun to explore the hypothesis that essential bacterial components, such as lipopolysaccharides (LPS) and peptidoglycan (PGN), which are well-characterized ligands for receptors expressed by immune cells, might also modulate host neuronal activity. Such molecular dialogue could allow the host to “sample” its surroundings via its sensory system and adopt behaviors that limit contact with pathogenic microbes or, in case of infection, reduce its impact on the host and its offspring (Kobler et al., 2020; Yanagawa et al., 2017). For instance, PGN detection by a few specific neurons can modulate oviposition behavior in infected females (Kurz et al., 2017; Masuzzo et al., 2022). In this case, the detection of a universal bacterial cell wall component by certain neurons in the brain acts as a molecular alarm, triggering a behavioral change in infected females, reducing their oviposition rate, and potentially conserving energy for the energy-demanding immune response. Other studies demonstrate that bacterial molecular patterns are directly detected by the fly’s sensory system. The detection of LPS by bitter taste neurons enables flies to avoid LPS in feeding and oviposition assays (Soldano et al., 2016). We previously reported that by activating the bitter gustatory network, bacterial PGN can suppress the appetitive effect of a sucrose solution, demonstrating that PGN can be perceived as a bitter substance by flies (Montanari et al., 2024).

In this study, we used the proboscis extension reflex assay (Shiraiwa and Carlson, 2007) and calcium imaging to assess whether a PGN solution could also activate the sweet taste circuit in flies. Our data reveal that PGN can elicit Gr5a+ neuron-dependent activation within the sweet taste network, demonstrating a dual response of the fly’s taste system to PGN. This response is repulsive at low concentrations and attractive at higher concentrations, differing from the dual response to salt or hexanoic acid observed in other gustatory neurons (Jaeger et al., 2018; McDowell et al., 2022; Pradhan et al., 2023; Zhang et al., 2013). This dichotomous response suggests that *Drosophila* shares a need to detect and differentiate between high and low concentrations of specific compounds present in food substrates, engaging different receptors and neuronal cells with distinct coding logic. Furthermore, the detection of bacteria by flies requires integrating inputs from multiple subsets of sensory neurons to generate appropriate responses to bacterial PGN.

## Results

### Bacterial PGN is Attractive to Flies at High Concentrations

We have previously shown that Diaminopimelic-type peptidoglycan (Dap-type PGN), which forms the cell wall of all Gram-negative bacteria (specifically *Escherichia coli* PGN in our study), activates neurons responsible for aversive behaviors in flies (Montanari et al., 2024). Specifically, Gr66a+ (Wang et al., 2004) and ppk23+ (Jaeger et al., 2018) adult neurons located on the proboscis respond to PGN (Masuzzo et al., 2022; Montanari et al., 2024). Moreover, overexpression of IMD pathway components in Gr66a+ cells alters fly behavior (Masuzzo et al., 2022). Finally, using the Proboscis Extension Reflex (PER) assay, which quantifies gustatory reflexes in immobilized animals (Shiraiwa and Carlson, 2007), we showed that *E. coli* PGN can evoke an aversive response in flies (Montanari et al., 2024).

Given the expected presence of bacteria on the natural food source of *Drosophila melanogaster*, we wondered whether bacteria might also be perceived as beneficial by the adult fly. To this end, we investigated the potential attractiveness of PGN. To assess whether brief contact with PGN could trigger an attractive response, we focused on the PER (Figure 1A) (Schwarz et al., 2017; Shiraiwa and Carlson, 2007). To accurately assess this response, we tested the molecule without mixing it with anything other than its solvent. When the fly’s labellum (analogous to the tongue) is stimulated by a molecule such as sucrose, it immediately extends its proboscis towards the food (Video 1). A given chemical is therefore considered more or less attractive based on its response relative to the reference solvent and a prototypical attractive substance, such as sucrose.

**Figure 1.**
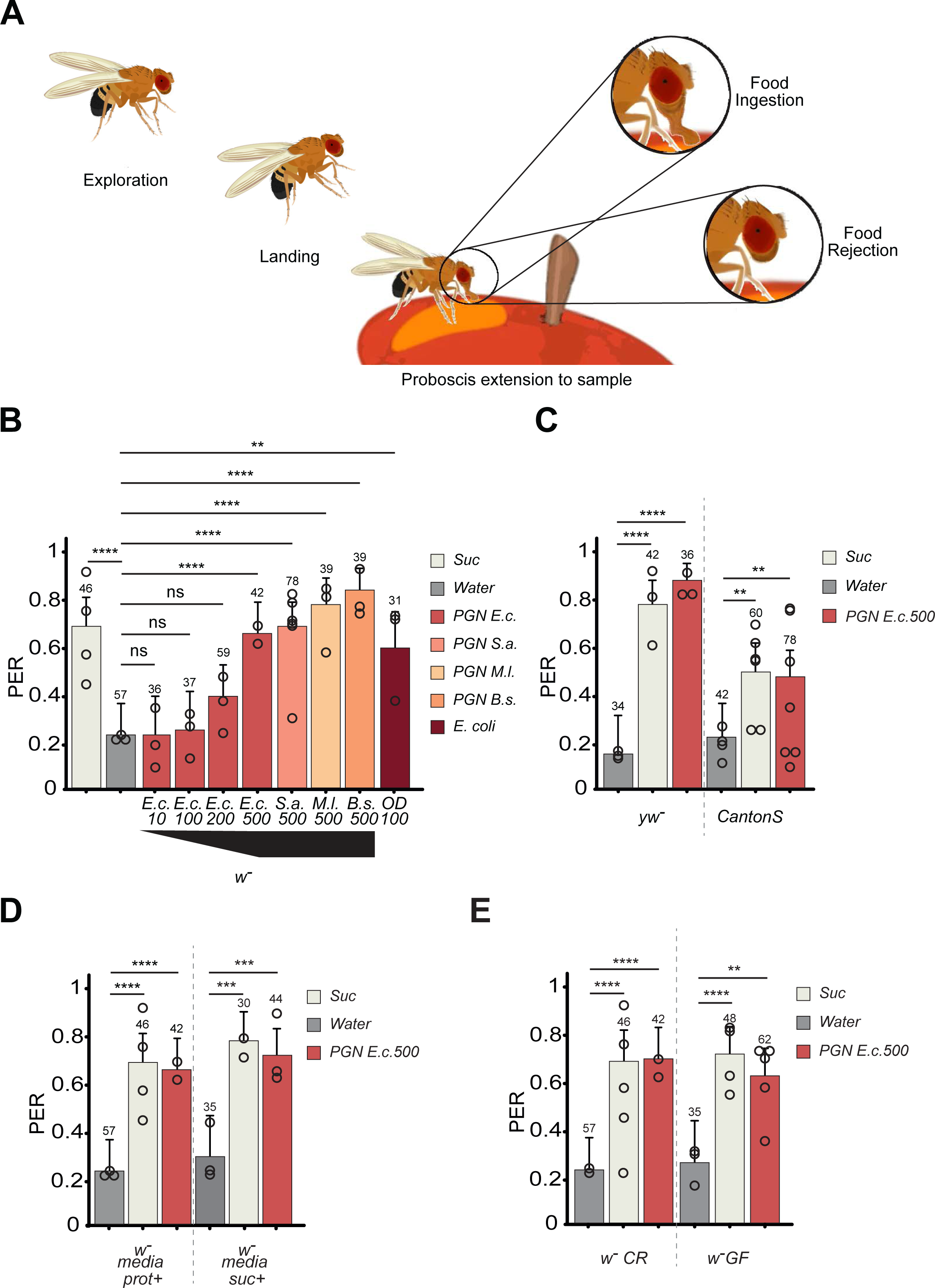
Dose-dependent attraction to PGN. (A) Drosophila feeding behavior is characterized by a sequence of distinct behavioral modules. A starved fly initiates a search for food, and the detection of food odors stops this search. The initial gustatory cues are acquired by sampling food using the proboscis. If the fly encounters a potentially appetizing stimulus, it will begin feeding; if the stimulus is aversive, the fly will retract its proboscis. (B) PER index of w− flies to control solutions of 1mM sucrose, water, and increasing concentrations of PGN from *E. coli* K12 (E.c.), *S. aureus* (S.a.), *M. luteus* (M.l.), and *B. subtilis* (B.s.), or an *E. coli* culture at OD100. The numbers below the x-axis represent the final PGN concentrations in µg/mL and the OD value for the *E. coli* culture. (C) Attraction to PGN is independent of genetic background. PER index of yw− and CantonS flies to control solutions of water, 1mM sucrose, and PGN from *E. coli* K12 at 500µg/mL. (D) Raising medium has no effect on attraction to PGN. PER index of w− flies raised on protein-enriched (prot+) or sucrose-enriched (suc+) media, to control solutions of water, 1mM sucrose, and PGN from *E. coli* K12 at 500µg/mL. Data for w− prot+ flies are the same as in panel 1A. (E) Germ-free status does not affect attraction to PGN. PER index of w− germ-free (GF) and conventionally raised (CR) flies to control solutions of water, 1mM sucrose, and PGN from *E. coli* K12 at 500µg/mL. Data for w− CR flies are the same as in panel 1A. The PER index is calculated as the percentage of flies that responded with a PER to the stimulation ± 95% confidence interval (CI). A PER value of 1 indicates that 100% of the tested flies extended their proboscis in response to the stimulus, while a value of 0.2 indicates that 20% of the flies responded. The number of flies tested (n) is shown above each bar. For each condition, at least three groups of 10 flies each were tested. “ns” indicates p>0.05, * indicates p<0.05, ** indicates p<0.01, *** indicates p<0.001, **** indicates p<0.0001, Fisher Exact Test. Further details are available in the figure section on lines, conditions, and statistics.

To determine if PGN could be attractive to flies, we tested this bacterial molecule alone, diluted in water at various concentrations and compared to sucrose. When the labellum was exposed to 1 mM sucrose, approximately 70% of flies extended their proboscis in an attempt to feed, illustrating the appetitive value of sucrose (Video 1 and Figure 1B). Water alone, the solvent for sucrose and PGN, triggered a PER in 20% of flies, and since the animals were allowed to drink to satiety before the test, we considered this response to water to be neutral, likely due to mechanical stimulation (Video 2 and Figure 1B). We then tested the fly’s response to *E. coli* DAP-type PGN. At concentrations of 10 µg/ml (E.c.10) and 100 µg/ml (E.c.100), only 20% of flies extended their proboscis, not different from the response to water, suggesting that PGN is not palatable at these concentrations (Figure 1B). However, exposure to PGN at 200 µg/ml (*E.c.200*) showed a trend toward increased response, with around 40% of flies performing a PER (Figure 1B). At 500 µg/ml (*E.c.500*), 70% of flies extended their proboscis, a response not different to that seen with 1 mM sucrose (Video 3, Video 4 and Figure 1B). Thus, PGN from *E. coli* at 500 µg/ml is attractive to flies.

When DAP-type PGN from the Gram-positive bacterium *Bacillus subtilis* (*B.s.500*) and Lysine-type PGNs from *Staphylococcus aureus* (*S.a.500*), *Micrococcus luteus* (*M.l.500*), and *E. coli* at an OD of 100 (OD100) were used as elicitors, the response was comparable to that of sucrose or *E. coli* PGN at 500 µg/ml (Figure 1B). The attractive effect of *E. coli* PGN at 500 µg/ml (*E.c.500*) was observed in flies from different genetic backgrounds, including *w-*, *yw*, and CantonS strains, as well as in *w-* flies reared on either protein-rich (prot+) or sugar-rich (suc+) media, demonstrating the robustness of the phenotype (Figure 1C and 1D).

These findings indicate that, at concentrations of 500µg/mL, both Lysine- and DAP-type PGNs are attractive to the fly’s gustatory system.

### Axenic flies are still capable of triggering the PER response to PGN

Given that both larvae and adults live in a non-sterile environment, and that we recently reported that larval cohabitation with bacteria is crucial for adults to perceive DAP-type PGN as aversive, we tested whether the palatability of PGN *E.c.500* in adult flies was also dependent on the presence of larval microbiota (Montanari et al., 2024). When flies were reared on antibiotic-containing medium throughout their lifespan (axenic), the PER response to DAP-type PGN *E.c.500* solutions was not different to that observed with sucrose and indistinguishable from the response measured in non-axenic animals (Figure 1E). Thus, unlike our previous findings regarding aversion to DAP-type PGN (Montanari et al., 2024), neither larval cohabitation with bacteria nor adult life in a non-aseptic medium influenced attractive behavior toward high concentrations of DAP-type PGN.

### PGN Detection by the Gustatory System Involves Gr5a Neurons

To identify which neurons respond to PGN E.c.500, attractive proboscis extension reflexes (PER) were performed in flies where specific neurons were inactivated by overexpressing the Kir2.1 potassium channel (UAS-Kir2.1), which activity impairs the generation of action potentials. The expression of the Gr5a receptor is characteristic of neurons involved in the detection of sweet molecules (Figure 2A) (Dahanukar et al., 2001), and these Gr5a+ neurons play a role in attractive behaviors toward sucrose or threalose. Importantly, inactivation of Gr5a+ neurons using the cell-specific driver Gr5a-Gal4 (Gr5a-Gal4/UAS-Kir2.1) abolished the attractive response to both 1mM sucrose and DAP-type PGN (Figure 2B). Thus, sweet neurons are essential for triggering a PER response to high concentrations of PGN in adult flies. Notably, the PER response of Gr5a-Gal4/UAS-Kir2.1 flies to PGN E.c.500 was significantly lower than the baseline response to water, with less than 10% of flies showing PER (Figure 2B). This suggests that in the absence of sweet neuron activity (Gr5a-Gal4/UAS-Kir2.1), PGN E.c.500 is perceived as aversive by the flies. This finding was consistent with our previous observation that DAP-type PGNs can induce ppk23+ neuron-dependent aversion (Montanari et al., 2024). A possible explanation is that, in the absence of attractive PGN perception (Gr5a-Gal4/UAS-Kir2.1), the aversion to this molecule becomes apparent. These results suggest an integration of positive and negative inputs that influence the fly’s decision-making process regarding the taste of different DAP-type PGN concentrations.

**Figure 2.**
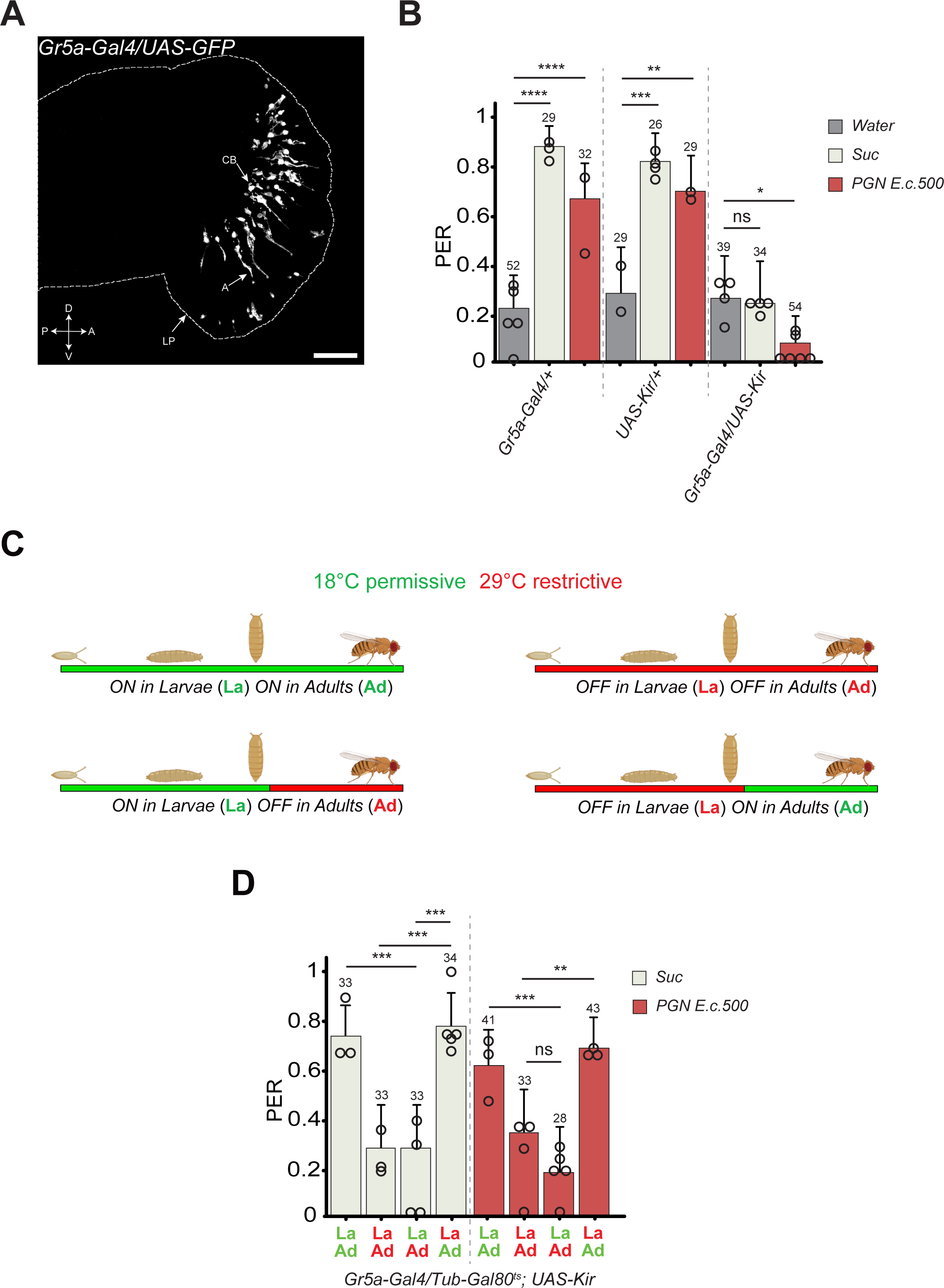
Gr5a+ neurons are required for attraction to PGN. (A) Confocal image of the proboscis of Gr5a-Gal4/UAS-GFP flies showing gustatory neurons in the fly proboscis. LP refers to Labellum-Proboscis, CB to cellular body; A denotes axons emanating from neurons in the proboscis. D, V, P, and A stand for Dorsal, Ventral, Posterior, and Anterior. (B) Impairing the activity of Gr5a+ neurons via UAS-Kir2.1 eliminates the attraction to PGN. PER index of flies to control solutions of water, 1mM sucrose, and PGN from *E. coli* at 500µg/mL. (C) Graph showing the life periods during which flies are shifted from 18°C (green) to 29°C (red). (D) Gr5a+ neurons are functionally required for PGN attraction in adults but are not needed during the larval stage. PER index of flies to 1mM sucrose and PGN from *E. coli* K12 at 500µg/mL. The ubiquitously expressed Tub-Gal80ts inhibits the activity of Gal4 at 18°C and is inactivated at 29°C, allowing the expression of UAS-Kir2.1 and the inhibition of Gr5a+ neuron activity. The PER index is calculated as the percentage of flies responding with a PER to the stimulus ± 95% CI. The number of flies tested (n) is shown above each bar. “ns” indicates p>0.05, * indicates p<0.05, ** indicates p<0.01, *** indicates p<0.001, **** indicates p<0.0001, Fisher Exact Test. Further details on conditions and statistics are provided in the figure section.

### Gr5a Neurons Are Required in Adults to Detect PGN E.c.500

In a previous study, we showed that Gr66a+ larval neurons are essential for priming the adult behavioral response to DAP-type PGN (Montanari et al., 2024). Despite the fact that Gr5a+ neurons do not seem to be present in larvae (Kwon et al., 2011), we wished to test in an unbiased way whether the activity of the larval cells we may target using Gr5a-Gal4 could influence adult attraction to PGN E.c.500. We used the Gal4/Gal80ts binary system, which allows spatial and temporal control of neuron inactivation when coupled with UAS-Kir2.1 (Figure 2C). When 1mM sucrose or PGN E.c.500 solutions were used as stimuli, conditional inactivation of Gr5a+ neurons only in adults was as effective as lifelong inactivation of these neurons. In contrast, inactivating Gr5a+ cells only during the larval stage had no impact on the fly’s ability to respond to sucrose or PGN E.c.500 (Figure 2D). Therefore, adult PER behavior towards PGN E.c.500 requires functional Gr5a+ neurons in the adult stage.

### DAP-type PGN Directly Activates Gr5a Sweet Neurons

The observation that Gr5a+ neurons are required for the PER response to PGN stimulation led us to investigate whether these cells could be directly activated by DAP-type PGN. Using *in vivo* GCaMP imaging, we monitored the neuronal activity of Gr5a+ axon projections in the subesophageal zone (SEZ) of the brain of Gr5a-Gal4/UAS-GCaMP6s flies, which were exposed to either water (Video 5) or a drop of PGN (Video 6), followed by 100mM sucrose (Video 7) to assess the viability and identity of neurons. Exposure of the labellum of *Gr5a-Gal4/UAS-GCaMP6s* flies to DAP-type PGN at 500µg/ml resulted in an increase in intracellular calcium levels in axonal projections in the SEZ (Figure 3A). These results demonstrate that PGN at high concentrations can directly activate sweet taste neurons.

**Figure 3.**
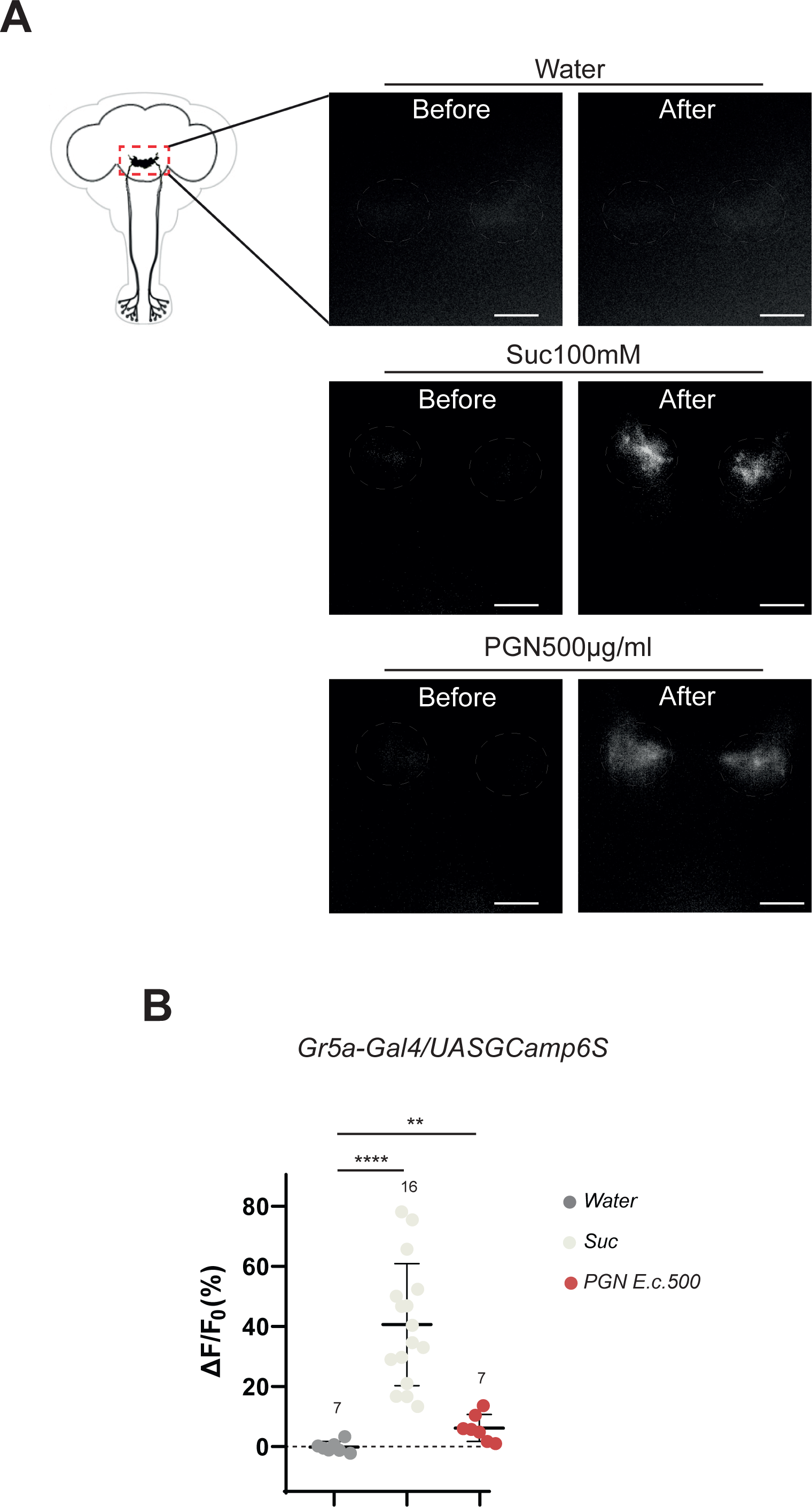
Real-time calcium imaging with GCaMP6s reflects the *in vivo* neuronal activity of Gr5a+ neurons. (A) Representative images showing the GCaMP6s intensity before and after the addition of control water, 100mM sucrose, or PGN from *E. coli* K12 at 500µg/mL. Scale bar is 10µm. (B) Averaged fluorescence intensity of negative peaks ± SEM for Gr5a+ neurons in response to water (n=7), 100mM sucrose (n=16), or PGN from *E. coli* K12 at 500µg/mL (n=7). In (A), ** indicates p<0.01, **** indicates p<0.0001, non-parametric t-test, Mann-Whitney test. Further details on conditions and statistics can be found in the figure section.

### Sweet Neurons Mediating Attraction to PGN E.c.500 Require Gr5a and Gr64 Receptors but Are Independent of the IMD Pathway

Having identified the neurons that respond to PGN and are required for the PER, we next investigated the receptors and pathways involved. Previous studies have shown that the Gr5a and Gr64a-f family proteins are receptors expressed in sweet neurons and are involved in detecting sweet molecules such as sucrose and fructose (Dahanukar et al., 2001; Jiao et al., 2008). To identify the receptors involved in PGN sensing, we used mutants lacking both Gr5a and the Gr64a-f family receptors (Gr5a-Gr64a-fCRISPR). The PER response of these mutants to sucrose and PGN E.c.500 was comparable to the response to water (Figure 4A). Since these mutant animals no longer respond to sweet molecules, we used a low salt concentration (50mM NaCl) as a positive attractive control to ensure proper selection of responsive animals. Of note, these mutant animals showed an increasing tendency towards an attraction to water and as anticipated, their response to sucrose was lower compared to wild-type insects and indistinguishable from the water baseline, confirming the role of these receptors. Importantly, the lack of increased PER upon exposure to PGN E.c.500, compared to the baseline water response, indicates that the tested sugar receptors are involved in PGN E.c.500 perception. Regarding the immune response, DAP-type PGN has been shown to be a major inducer of the PGRP/NF-kB-IMD pathway, leading to the production of immune effectors and regulators (Leulier et al., 2003) (Figure 4B). Our previous data indicated that elements of the IMD pathway are required for DAP-type PGN signal transduction in adult bitter taste neurons (Gr66a+ cells) (Masuzzo et al., 2022). To explore the role of the IMD pathway in PGN E.c.500 response, we used RNA interference to downregulate components of the IMD pathway. Flies with downregulated PGRP-LC receptor and the IMD transducer FADD specifically in Gr5a+ neurons (*Gr5a-Gal4/UAS-pgrp-LC_RNAi* and *Gr5a-Gal4/UAS-Fadd_RNAi*) showed sensitivity to PGN E.c.500 similar to that of control flies, suggesting that contrary to our previous findings regarding Gr66a+ neurons and PGN in adults (Masuzzo et al., 2022), neither PGRP-LC nor FADD is required in Gr5a+ neurons for the PER response to PGN E.c.500 (Figure 4C).

**Figure 4.**
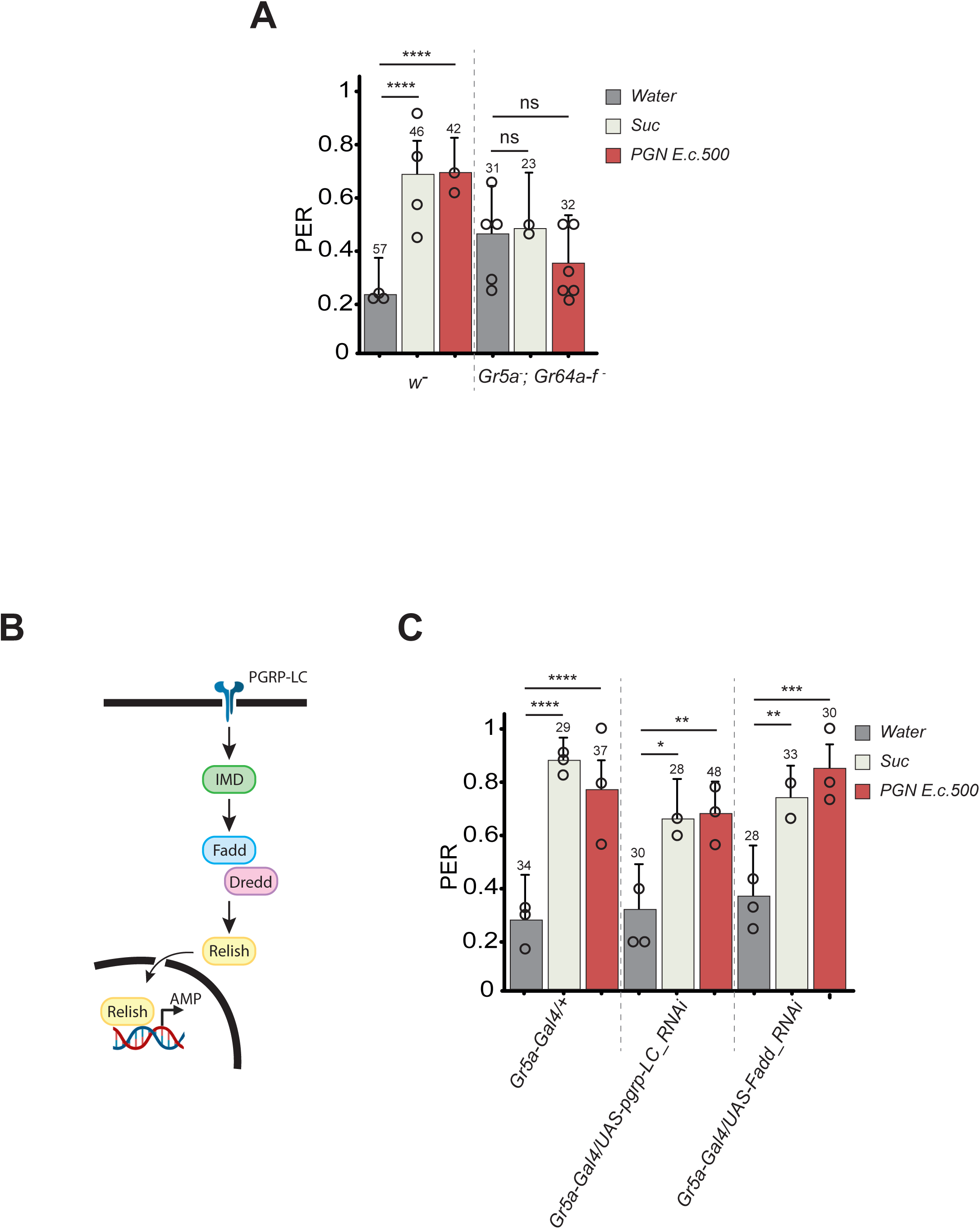
Sugar receptors are involved in PGN attraction. (A) PER index of mutant flies lacking the Gr5a receptor as well as the Gr64a-f family to water, 1mM sucrose, and PGN from *E. coli* K12 at 500µg/mL, compared to w− flies. Data for w− flies are the same as in Fig. 1A. (B) Diagram of the IMD pathway. (C) RNAi-mediated inactivation of PGRP-LC (UAS-pgrp-LC RNAi) or Fadd (UAS-Fadd RNAi) in Gr5a+ cells does not affect attraction to PGN. PER index of flies to water, 1mM sucrose, and PGN from *E. coli* K12 at 500µg/mL. For (A) and (C), the PER index is calculated as the percentage of flies responding with a PER to the stimulus ± 95% CI. The number of flies tested (n) is shown above each bar. “ns” indicates p>0.05, * indicates p<0.05, ** indicates p<0.01, *** indicates p<0.001, **** indicates p<0.0001, Fisher Exact Test. Further details on conditions and statistics are available in the figure section.

### Integration of Attractive and Aversive Signals from PGN Exposure Modulates the PER

Our data show that PGN E.c.500 elicits a Gr5a+ neuron-dependent PER response, comparable to that induced by 1mM sucrose. We also demonstrated that DAP-type PGN at concentration of 500µg/ml is capable of activating Gr5a+ neurons (Figure 3A). We previously reported that DAP-type PGN at 200µg/ml antagonizes the PER response to 1mM sucrose by activating adult ppk23+ neurons (Montanari et al., 2024). Taken together, these data suggest that DAP-type PGN can trigger behaviors involving both attractive and aversive neurons. Thus, it is possible that depending on its concentration, *E. coli*-derived DAP-type PGN can either trigger or inhibit a PER via the activation of two independent sets of neurons (Figure 5A).

**Figure 5.**
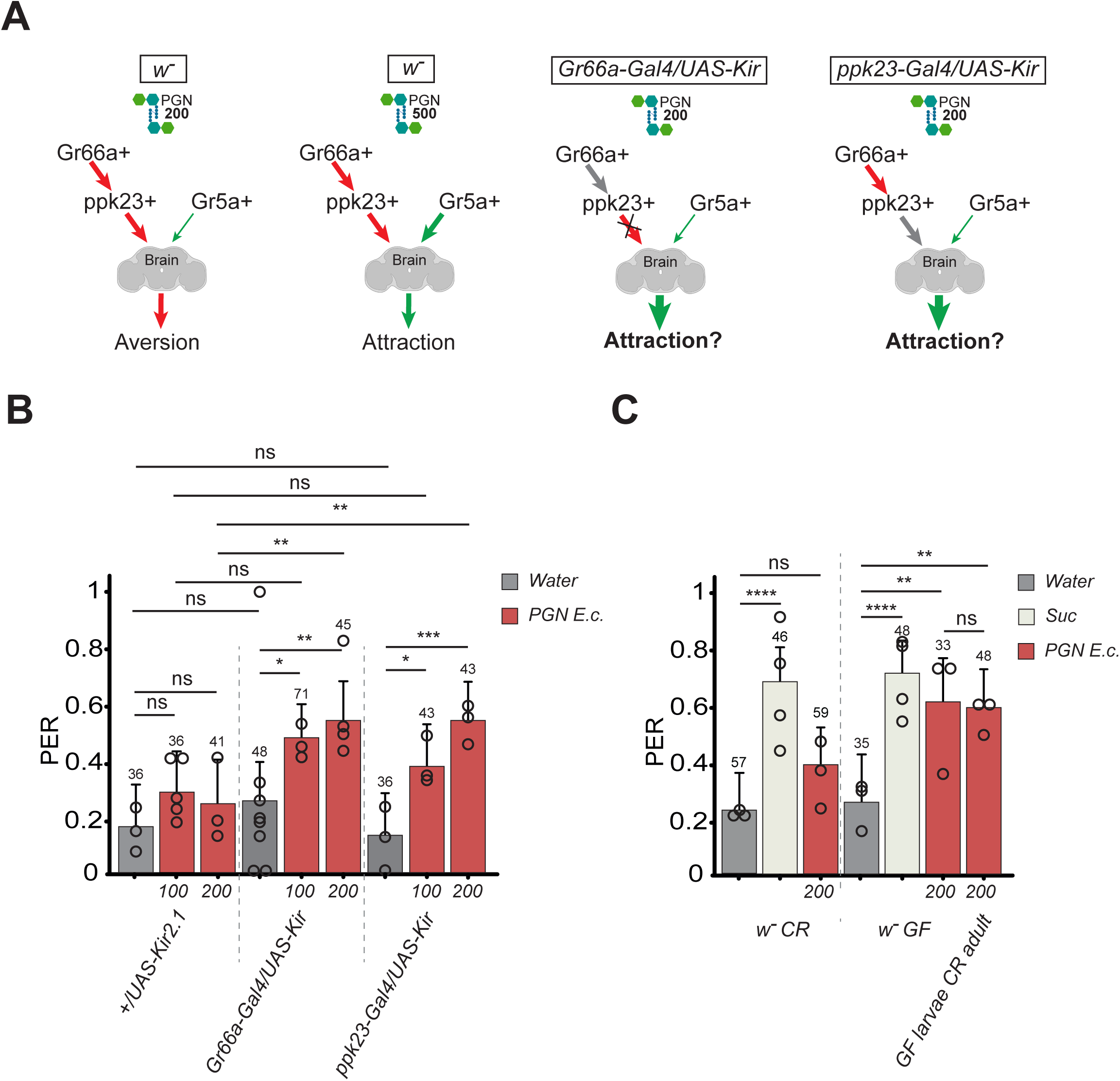
Integration of Attractive and Aversive Signals from PGN Exposure Modulates the PER. (A) Schematic illustrating the balance between attraction and aversion to PGN and the effects of genetic manipulations in Gr66a+ and ppk23 neurons on this balance. Our previous work showed that Gr66a+ neurons are not required during adulthood for PGN aversion, but their activity during the larval stage is necessary for ppk23+ neurons to recognize and show aversion to PGN in adulthood (arrow linking Gr66a+ to ppk23+). At a PGN concentration of 500µg/mL, input from Gr5a+ neurons shifts the balance from aversion to attraction. Impairing Gr66a+ or ppk23+ neuron activity (Gr66a-Gal4/UAS-Kir2.1 and ppk23-Gal4/UAS-Kir2.1) abolishes their input, leading to increased attraction to PGN. (B) Impairing the activity of Gr66a+ or ppk23+ neurons with UAS-Kir2.1 increases attraction to PGN at 200µg/mL. PER index of flies to control solutions of water and PGN from *E. coli* K12 at 100µg/mL or 200µg/mL. (C) PER index of flies raised as conventional (CR), germ-free (GF), or germ-free only during the larval stage (GF larvae CR adult) to control solutions of water and PGN from *E. coli* K12 at 200µg/mL. Data for w− CR flies are the same as in Fig. 1A. For (B) and (C), the PER index is calculated as the percentage of flies responding with a PER to the stimulus ± 95% CI. The number of flies tested (n) is shown above each bar. “ns” indicates p>0.05, * indicates p<0.05, ** indicates p<0.01, *** indicates p<0.001, **** indicates p<0.0001, Fisher Exact Test. Further details on conditions and statistics are available in the figure section.

Therefore, when exposed to PGN, opposing signals from different neurons can be integrated by the fly, and the resulting PER will be the outcome of this integration. In this context, the PER to PGN E.c.500 is likely the sum of inputs from Gr5a+ neurons (attraction) as well as from Gr66a+ or ppk23+ neurons (aversion). In the context of a PGN concentration triggering attraction, blocking the input from sweet neurons using Gr5a-Gal4/UAS-Kir2.1 should abolish the attraction and in turn reveal an aversion. Intriguingly, the PER to PGN E.c.500 was lower than the water baseline in Gr5a-Gal4/UAS-Kir2.1 flies indicating a loss of attraction and suggesting an aversion (Figure 2B). This observation was further confirmed by performing the opposite experiment, where blocking the activity of aversive neurons should lead to an increase in attraction. Indeed, while the attractive PER to DAP-type PGN at 100 µg/mL or 200 µg/mL in wild-type flies was not statistically different from the solvent control (Figure 1B and 5B), blocking the activity of Gr66a+ or ppk23+ neurons (Gr66a-Gal4/UAS-Kir2.1 and ppk23-Gal4/UAS-Kir2.1) resulted in a significantly higher PER to these PGN concentrations compared to the water baseline, concentrations that do not elicit attraction in standard conditions (Figure 5A and 5B).

We previously reported that aversion to PGN requires the activity of adult ppk23+ neurons, and that these neurons’ ability to respond to PGN is dependent on larval cohabitation with bacteria (Montanari et al., 2024). Based on this, we hypothesized that rearing animals in axenic conditions would impair PGN aversion in adults, making them more attracted to lower PGN concentrations that do not elicit attraction in standard conditions. While the PER to water or 1mM sucrose was not influenced by rearing conditions, axenic animals exposed to DAP-type PGN at 200µg/ml exhibited an increased PER compared to controls, regardless of whether the axenic period lasted throughout their lives or was restricted to the larval stage (Figure 5C). This result supports a model in which the PER to PGN is determined by an integration of attractive and aversive signals from independent neuronal networks.

## Discussion

Our results demonstrate that at concentrations above 200 µg/ml, both DAP-type and Lys-type PGN, essentially the majority of PGNs, are capable of stimulating sweet neurons through the activation of receptors previously identified as sugar receptors. This activation might be due to the sugar components of PGN, specifically the long polymers of the disaccharides N-acetylmuramic acid and N-acetylglucosamine. Since these sugar components are common to all PGNs, it may explain why both Lys- and DAP-type PGNs are able to trigger attraction, as observed in this study (Vollmer et al., 2008).

PGN can be modified by enzymes like amidases and lysozymes, which fragment the polymer into smaller units called muropeptides. Previous studies in insects and mammals have shown that different muropeptides can activate specific receptors with varying degrees of efficacy (Fioriti et al., 2024; Leulier et al., 2003; Tosoni et al., 2019). The minimum structural unit of PGN capable of activating sugar neurons remains to be determined and whether the level of neuronal activation depends on the specific structure of the muropeptides.

Interestingly, the activation we observed appears to be independent of the IMD pathway, as the proboscis extension responses (PER) to PGN were similar in both control flies and those RNAi inactivated for certain components of the IMD/NF-kB pathway. This response contrasts with previous findings where PGN, at concentrations near 200 µg/ml, was shown to activate adult bitter Gr66a neurons via part of the IMD pathway, though not involving the Relish transcription factor (Masuzzo et al., 2022). It is also worth noting that the ability to perceive PGN as a bitter molecule has only been observed with DAP-type PGNs, which are typically found in Gram-negative bacteria, making this response specific to a subset of bacterial species, unlike the response observed in our study.

In contrast, the sweet neuron response we report here more closely resembles the response observed in ppk23+ neurons, which is also independent of the IMD pathway (Montanari et al., 2024). However, there is a notable difference between the attractive response mediated by Gr5a+ neurons and the aversive response mediated by ppk23+ neurons. The aversive response to PGN requires bacterial priming during the larval phase and is absent in axenic flies, while the Gr5a+ neuron-mediated attraction to PGN occurs even in flies obtained from larva reared in the complete absence of bacteria. These findings suggest that the detection and response to PGN by Gr5a+ neurons in adults are not dependent on prior exposure to bacteria.

By measuring PERs in flies with blocked bitter or sweet circuits, we show that the fly’s taste system integrates signals from at least three distinct types of neurons—Gr66a+, ppk23+, and Gr5a+—to determine how to respond to PGN. Notably, the behavior requiring these neurons is not triggered by the same concentrations of PGN. Similar dose-dependent behavioral responses have been described for other compounds. For example, NaCl is attractive at 50 mM through the Gr5a+ network, but aversive at 250 mM via ppk23+ neurons (Zhang et al., 2013; Jaeger et al., 2018; McDowell et al., 2022). Likewise, hexanoic acid is attractive at low concentrations via Gr5a+ cells, but aversive at higher concentrations through the bitter network (Pradhan et al., 2023). These examples support the idea that different concentrations of the same compound can trigger opposite behaviors, depending on the involved neuronal circuits.

Some studies suggest a more complex scenario, where a molecule can be attractive as an odor but aversive as a tastant. A prime example of this is acetic acid, which is attractive when detected by the gustatory system but repulsive when sensed by the olfactory system (Devineni et al., 2019; Joseph et al., 2009). Additionally, systems that integrate opposite inputs from both sweet and bitter signals have been identified, such as the inhibition of sweet neurons by bitter neurons through GABA signaling (Chu et al., 2014) or by the OBP49a protein, which attenuates sweet neuron firing when bitter chemicals are combined (Jeong et al., 2013). These mechanisms help facilitate the avoidance of harmful compounds when sweet molecules are present in mixtures. In our case, the concentration that triggers aversion (200 µg/ml) is lower than the one that triggers attraction (500 µg/ml), suggesting a unique pattern of integration for PGN.

While our data go against the intuitive idea that the more you sense PGN, the more you should avoid it, one hypothesis could be that more bacteria equal an advanced decaying fruit with potentially more nutrients. Few bacteria may indicate the presence of a not so advanced decaying fruit. Importantly, we need to consider that the adult aversion to PGN is related to the exposure to pathobionts during the larval life (Montanari et al., 2024). If a larva is only exposed to symbionts, the effect of PGN over adults will be as depicted with the adults of Figures 4B and 4C, with attractions to PGN concentrations as low as 100µg/ml. Thus, the naive adult (Germ-free raised) seems to be primarily tuned toward attraction to lower PGN concentrations, but this can be modulated in the context of a larvae that experienced a stress related to a bacteria like *L. brevis*. Indeed, our findings using the absence of the aversive response (i.e., in Gr66a-Gal4/UAS-Kir2.1 flies) highlight the fact that PGN concentrations at 100 µg/ml can trigger attraction. This suggests that sweet neurons are ‘naturally’ tuned to PGN concentrations lower than those that elicit aversion and lower than those that elicit attraction in ‘non-naïve’ animals. Taken together, it appears that the integration of bitter signals overrides the attraction, and the PER response toward PGN will necessitate a higher PGN concentration, as if the nutritional value of the environment must outweigh the risk associated with the presence of bacteria, similar to a benefit/risk balance. In a more complex, physiological context where both sweet and bitter molecules are present, the overall influence of microbial molecules like PGN may be balanced by the other chemicals present.

Finally, it remains to be explored what these PGN concentrations represent in terms of bacterial populations in natural environments, and whether the physiological state of the bacteria influences their ability to release PGN and if so, in which forms and at what concentrations. This information would be critical for understanding the role of PGN detection systems in nature.

This task is further complicated by the fact that PGN perception is just one small piece of the sensory information that flies use to navigate and make decisions in an environment teeming with bacteria and other microorganisms. This complex sensory integration is key to the fly’s ability to balance the need for food, reproduction, and immunity in an ever-changing microbial landscape.

## Acknowledgments

This work was supported by CNRS, ANR BACNEURODRO (ANR-17-CE16-0023-01), Equipe Fondation pour la Recherche Médicale (EQU201603007783) et l’Institut Universitaire de France to J.R. and the ANR Pepneuron (ANR-21-CE16-0027) to J.R. and Y.G. Research in Y.G.’s laboratory is supported by the CNRS, the “Université de Bourgogne Franche-Comté”, the Conseil Régional Bourgogne Franche-Comté (PARI grant), the FEDER (European Funding for Regional Economical Development), and the European Council (ERC starting grant, GliSFCo-311403)

## Material and methods

### Fly husbandry

Flies were grown at 25°C on a yeast/cornmeal medium in 12h/12h light/dark cycle-controlled incubators. For 1 L of food, 8.2 g of agar (VWR, cat. #20768.361), 80 g of cornmeal flour (Westhove, Farigel maize H1) and 80 g of yeast extract (VWR, cat. #24979.413) were cooked for 10 min in boiling water. 5.2 g of Methylparaben sodium salt (MERCK, cat. #106756) and 4 mL of 99% propionic acid (CARLOERBA, cat. #409553) were added when the food had cooled down (prot+ media).

suc+ media: for 1 L of food, 11 g of agar (VWR, cat. #20768.361), 80 g of cornmeal flour (Westhove, Farigel maize H1), 20 g of yeast extract (VWR, cat. #24979.413) and 30g of Sucrose were cooked for 10 min in boiling water. 2.5g of Moldex and 5 mL of 99% propionic acid (CARLOERBA, cat. #409553) were added when the food had cooled down.

### Fly stocks

The reference strain in this study is noted *w^−^* and corresponds to *w^[1118]^* (Bloomington #5905). Canton S (Bloomington #64349) and *y^[1]^, w^[1118]^*and *Gr5A^−^; Gr64a-f^−^* (gift from Bernard Charroux) and *Gr5a-Gal4* (Bloomington #57992) and *UAS-GCaMP6S* (Bloomington #42746) and *UAS-Kir2.1* (Bloomington #6595) and *Tub-Gal80ts* (Bloomington #7016) and *UAS-GFP* (Bloomington #32195) and *UAS-pgrp-LC_RNAi* (VDRC #101636) and *UAS-Fadd_RNAi* (Leulier et al., 2002) and *ppk23-Gal4* (Bloomington #93026) and *Gr66a-Gal4* (Bloomington #28801).

### PER assay

All flies used for the test were females between 5 and 7 days old. Unless experimental conditions require it, the flies are kept and staged at 25°C to avoid any temperature changes once they are put into starvation. The day before, the tested flies are starved in an empty tube with water-soaked plug for 24h at 25°C.

Eighteen flies are tested in one assay, 6 flies are mounted on one slide and in pairs under each coverslip. To prepare the slide, three pieces of double-sided tape are regularly spaced on a slide. Two spacers are created on the sides of each piece of tape by shaping two thin cylinders of UHT paste. To avoid the use of carbon dioxide flies are anesthetized on ice. Under the microscope, two flies are stuck on their backs, side by side, on same piece of tape so that their wings adhere to the tape. A coverslip is then placed on top of the two flies and pressed onto the UHT paste, blocking their front legs and immobilizing them.

Once all slides are prepared, they are transferred to a humid chamber and kept at 25°C for 1.5 hours to allow the flies to recover before the assay.

Flies are tested in pairs, the test is carried out until completion on a pair of flies (under the same coverslip), and then move on to the next pair.

Before the test, water is given to each pair of flies to ensure that the flies are not thirsty and do not respond with a PER to the water in which the solutions are prepared. Stimulation with the test solution is always followed by a control stimulation with a sweet solution, to assess the fly’s condition and its suitability for the test. During the test small strips of filter paper are soaked in the test solution and used to contact the fly’s labellum (three consecutive times per control and test phase). Contact with the fly’s proboscis should be as gentle as possible. Ideally the head should not move. A stronger touch may prevent the fly from responding to subsequent stimulation. Based on the protocol needed the test is done following the sequence and the timing in the table below.

**Tab.1.**
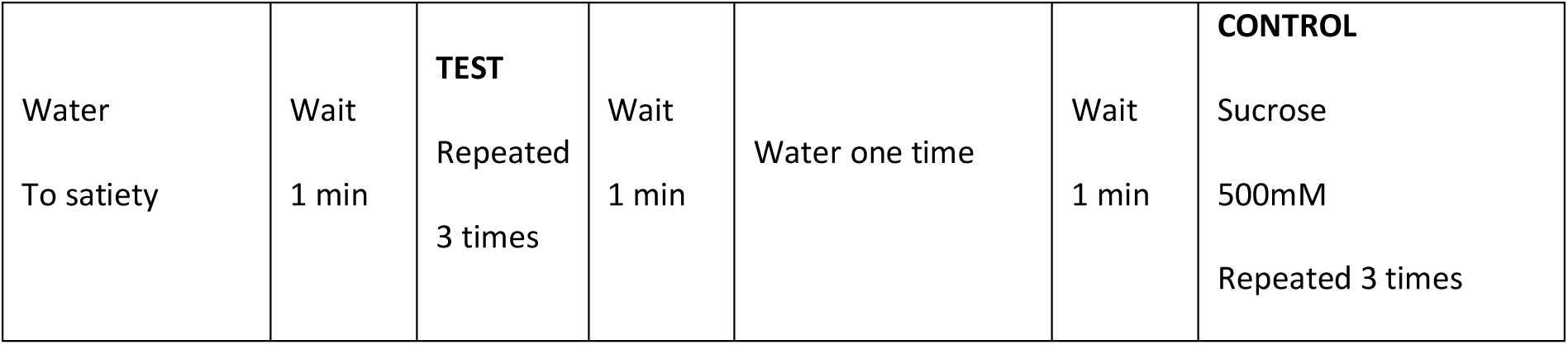
Attraction protocol.

All solutions to be tested are prepared the test day and stored at room temperature. The control stimulation at the end of the assay is performed with 500mM sucrose (D(+)-sucrose ≥99.5 %, p.a. Carl Roth GmbH + Co. KG). After each test or control stimulation, a water-soaked strip is used to tap the proboscis and clean it. For the water baseline, a paper stripe soaked in water is used as the test solution.

The response of the fly to each stimulation is recorded and averaged. A fly is considered as responding when a full extension of the proboscis is observed (PER). The duration of the extension is not taken into account. Flies that respond positively (PER) to at least one of the control stimulations are considered for further analysis, the others are discarded. The PER index is calculated as the ratio of flies tested that responded with a PER to the TEST stimulation (number of flies that responded at least 1 time to the test solution and at least 1 time to the control solution) / (total amount of flies that responded at least 1 time to the control solution) and represented as ± 95% CI. In case of stage dependent experiments, flies are shifted from one condition to another upon hatching.

### Microscopy

No immunostaining was performed with proboscises. Proboscises of adult females were dissected in PBS, rinsed with PBS, and directly mounted on slides using Vectashield fluorescent mounting medium. The tissues were visualized directly after. Images were captured with a LSM 780 Zeiss confocal microscope (20X air objective was used).

### *In vivo* calcium imaging

*In vivo* adult calcium imaging experiments were performed on 5-7 day-old starved mated females. Animals were raised on conventional media with males at 25°C. Flies were starved for 20-24 h in a tube containing a filter paper soaked in water prior to experiments. Flies of the appropriate genotype were anesthetized on ice for 1 h. Female flies were suspended by the neck on a plexiglass block (2 x 2 x 2.5 cm), with the proboscis facing the center of the block. Flies were immobilized using an insect pin (0.1 mm diameter) placed on the neck. The ends of the pin were fixed on the block with beeswax (Deiberit 502, Siladent, 1345 209212). The head was then glued on the block with a drop of rosin (Gum rosin, Sigma-Aldrich 1346 -60895-, dissolved in ethanol at 70 %) to avoid any movements. The anterior part of the head was thus oriented towards the objective of the microscope. Flies were placed in a humidified box for 1 h to allow the rosin to harden without damaging the living tissues. A plastic coverslip with a hole corresponding to the width of the space between the two eyes was placed on top of the head and fixed on the block with beeswax. The plastic coverslip was sealed on the cuticle with two-component silicon (Kwik-Sil, World Precision Instruments) leaving the proboscis exposed to the air. Ringer’s saline (130 mM NaCl, 5 mM KCl, 2 mM MgCl2, 2 mM CaCl2, 36 mM saccharose, 5 mM HEPES, pH 7.3) was placed on the head. The antenna area, air sacs, and the fat body was removed. The esophagus was cut without damaging the brain and taste nerves to allow visual access to the anterior ventral part of the sub-esophageal zone (SEZ). The exposed brain was rinsed twice with Ringer’s saline. GCaMP6s fluorescence was viewed with a Leica DM600B microscope under a 40x water objective. Stimulation was performed manually using a pipette with gel loading tip by applying 140 µL of tastant solution diluted in water on the proboscis. The gustatory stimulation was continuous as the tastant is in the drop contacting the proboscis. The recording started before the addition of the tastant and the calcium response could be observed immediately following the contact with the sensilla. For each condition, n=7 to 16. To ensure that GR5a neurons were still alive after application of water or of PGN at 500µg/ml, we used 100mM sucrose as a positive control. Consequently, for the sucrose, n = sum of flies tested with water + flies tested with PGN. Some quantifications with water or PGN were discarded due to fly movements, whereas they could be utilized for the sucrose.

GCaMP6s was excited using a Lumencor diode light source at 482 nm ± 25. Emitted light was collected through a 505-530 nm band-pass filter. Images were collected every 500ms using a Hamamatsu/HPF-ORCA Flash 4.0 camera and processed using Leica MM AF 2.2.9. Each experiment consisted of a recording of 70-100 images before stimulation and 160 images after stimulation. Data were analyzed as previously described (Silbering et al., 2012) by using FIJI (https://fiji.sc/).

### Different PGN used

*E. coli* K12: Invivogen, catalog code #tlrl-kipgn

*Staphylococcus aureus :* Invivogen, catalog code #tlrl-kipgn

*Micrococcus luteus* : Sigma Aldrich, catalog #53243

*Bacillus subtilis*: Sigma Aldrich, catalog #SMB00288

### Bacterial strains and maintenance

For this study, the following bacterial strain was used: *E. coli* K12 ATCC

Grown in LB liquid, in aerobic conditions, 37°C, shaking at 200 rpm.

### Statistics and data representation

The Prism software was used for statistical analyses.

For *in vivo* calcium imaging, the D’Agostino–Pearson test to assay whether the values are distributed normally was applied. As not all the data sets were considered normal, non-parametric statistical analysis such as non-parametric unpaired Mann–Whitney two-tailed tests was used for all the data presented.

For PER datasets. As the values obtained from one fly are categorical data with a *Yes* or *No* value, we used the Fisher exact t-test and the 95% confidence interval to test the statistical significance of a possible difference between a test sample and the related control.

For PER assays, at least 2 independent experiments were performed. The results from all the experiments were gathered and the total amount of flies tested is indicated in the graph. In addition, we do not show the average response from one experiment representative of the different biological replicates, but an average from all the data generated during the independent experiments in one graph. However, each open circle represents the average PER of 1 experiment.

### Detailed lines, conditions and statistics for the figure section

All these data are downloadable with the source data file.

DOI: 10.6084/m9.figshare.27908889

https://figshare.com/ndownloader/files/50813955

---

**Video 1 PER is triggered following exposure to sucrose 1mM**

https://doi.org/10.6084/m9.figshare.27929532.v1

Response following proboscis contact with a stripe of paper soaked in a 1mM solution (water as solvent). 7-days old females starved 15h prior to the assay.

**Video 2 PER is not triggered following exposure to wate**

https://doi.org/10.6084/m9.figshare.27929517.v1

Absence of response following proboscis contact with a stripe of paper soaked in water. 7-days old females starved 15h prior to the assay.

**Video 3 PER is triggered following exposure to PGN of *E. coli* at 500µg/ml**

https://doi.org/10.6084/m9.figshare.27929547.v1

Response following proboscis contact with a stripe of paper soaked in an *E. coli* PGN solution at 500µg/ml (water as solvent). 7-days old females starved 15h prior to the assay.

**Video 4 Comparison of Behavioral Response to Water and E. coli PGN in the Same Fly**

https://doi.org/10.6084/m9.figshare.27929553.v1

Using the same fly, this experiment compares the absence of response to a proboscis contact with a paper strip soaked in water, versus a clear response to a paper strip soaked in an *E. coli* PGN solution (500 µg/ml, with water as the solvent). The test was conducted on 7-day-old females, starved for 15 hours prior to the assay.

**Video 5 Gr5a+ neurons do not respond to water**

https://doi.org/10.6084/m9.figshare.28480388.v1

Real-time calcium imaging in the SEZ of adult females using the calcium indicator GCaMP6s under the control of the Gr5a-Gal4 driver to assess the in vivo neuronal activity of labellar Gr5a+ neurons following exposure to water. The water was added at t+ 2 seconds.

**Video 6 Gr5a+ neurons respond to DAP-type peptidoglycan**

https://doi.org/10.6084/m9.figshare.28479926.v1

Real-time calcium imaging in the SEZ of adult females using the calcium indicator GCaMP6s under the control of the Gr5a-Gal4 driver to assess the in vivo neuronal activity of labellar Gr5a+ neurons following exposure to *E. coli* PGN solution at 500µg/mL. The PGN was added at image 100 which corresponds to t+2 seconds.

**Video 7 Gr5a+ neurons respond to sucrose**

https://doi.org/10.6084/m9.figshare.28480460.v1

Real-time calcium imaging in the SEZ of adult females using the calcium indicator GCaMP6s under the control of the Gr5a-Gal4 driver to assess the in vivo neuronal activity of labellar Gr5a+ neurons following exposure to sucrose 100mM. The sucrose was added at image 80 which corresponds to t+2 seconds.

